# Monosomes actively translate synaptic mRNAs in neuronal processes

**DOI:** 10.1101/687475

**Authors:** Anne Biever, Caspar Glock, Georgi Tushev, Elena Ciirdaeva, Julian D. Langer, Erin M. Schuman

## Abstract

In order to deal with their huge volume and complex morphology, neurons localize mRNAs and ribosomes near synapses to produce proteins locally. A relative paucity of polyribosomes (considered the active sites of translation) detected in electron micrographs of neuronal processes (axons and dendrites), however, has suggested a rather limited capacity for local protein synthesis. Polysome profiling together with ribosome footprinting of microdissected synaptic regions revealed that a surprisingly high number of dendritic and/or axonal transcripts were predominantly associated with monosomes (single ribosomes). Contrary to prevailing views, the neuronal monosomes were in the process of active protein synthesis (e.g. they exhibited elongation). Most mRNAs showed a similar translational status in both compartments, but some transcripts exhibited differential ribosome occupancy in the somata and neuropil. Strikingly, monosome-preferred transcripts often encoded high-abundance synaptic proteins. This work suggests a significant contribution of monosome translation to the maintenance of the local neuronal proteome. This mode of translation can presumably solve some of restricted space issues (given the large size of polysomes) and also increase the diversity of proteins made from a limited number of ribosomes available in dendrites and axons.

## Main text

RNA sequencing and *in situ* hybridization have revealed the presence of an unexpectedly high number of RNA species in dendrites and/or axons of the CA1 neuropil (*1, 2*) and many studies have documented the local translation of proteins in dendrites and/or axons (3–5). During mRNA translation multiple ribosomes can occupy an individual mRNA (called a “polysome”) resulting in the generation of multiple copies of the encoded protein. Polysomes, usually recognized in electron micrographs as a cluster comprising 3 or more ribosomes, have been detected in neuronal dendrites (6, 7) but are surprisingly infrequent (e.g. < 0.5 polysome per µm, see (7)) given the diversity of mRNAs present in dendrites and axons (8). In neuronal processes, the features and mechanisms of translation have not been explored in detail, in part because of the relative inaccessibility of the relevant compartments (e.g. the dendrites and axons in the neuropil). In particular, how diverse proteins might be synthesized from a limited population of polysomes present in small synaptic volumes is an open question.

### Monosomes are the predominant ribosome population in neuronal processes

To visualize the capacity for protein synthesis in the neuropil *in vivo*, we labeled the *de novo* proteome using puromycylation (9, 10). We infused puromycin directly into the lateral ventricle of mice, waited 10 min, and then visualized newly synthesized proteins in hippocampal pyramidal neurons by co-immunofluorescence labeling of nascent protein (anti-puromycin antibody) and CA1 pyramidal neurons (anti-wolframin antibody; Wfs1). As expected, we detected an intense nascent protein signal in the somata layer (*stratum pyramidale*), comprising the cell bodies of pyramidal neurons (Fig. 1A, fig. S1). There was also strong nascent protein evident throughout the dendrites of pyramidal neurons in the neuropil (*stratum radiatum*) (Fig. 1A, fig. S1). Co-injection of a protein synthesis inhibitor (anisomycin) abolished the nascent protein signal. Because of the very short window of metabolic labeling, these data indicate that protein synthesis also occurs in dendrites *in vivo*.

**Fig. 1.**
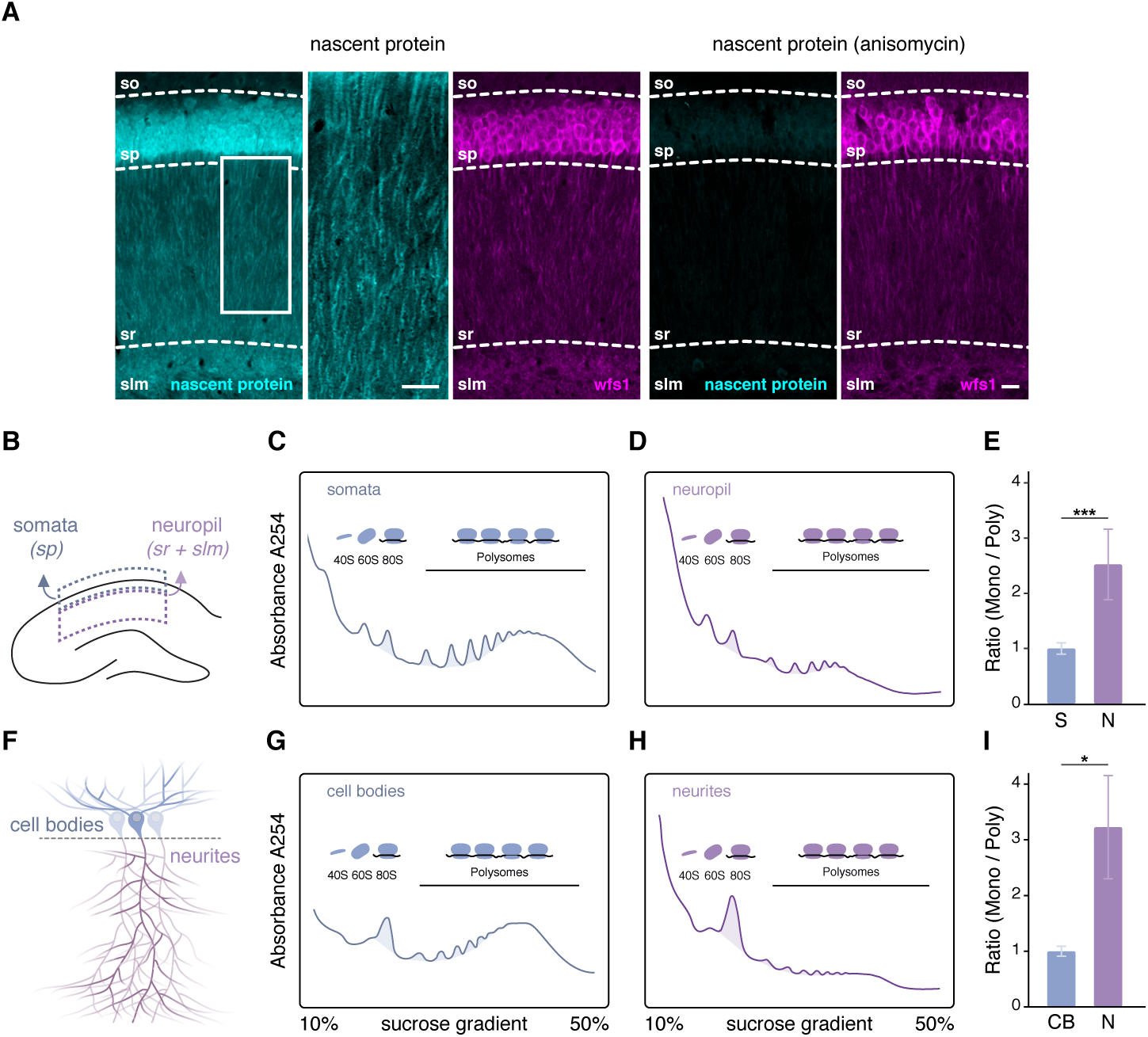
Monosomes are the major ribosome population in neuronal processes. **(A)** Immunofluorescence labeling of the nascent protein metabolic label (cyan) and the *Cornu Ammonis* 1 (CA1) pyramidal neuron marker Wfs1 (purple) in hippocampal sections from mice that received a brief infusion of puromycin without (left) or with the protein synthesis inhibitor anisomycin (right) into the lateral ventricle. Scale bar = 20 µm. A higher magnification image of the nascent protein signal in the boxed dendritic region is shown. Scale bar = 50 µm. so, *stratum oriens*; sp, *stratum pyramidale*; sr, *stratum radiatum*, slm, *stratum lacunosum moleculare*. **(B)** Scheme of a hippocampal slice showing the regions (somata and neuropil) that were microdissected for subsequent polysome profiling. Representative polysome profiles **(C and D)** and monosome to polysome (Mono/Poly) ratios **(E)** of the microdissected somata (blue) or neuropil (purple) (n= 7 biological replicates). Areas measured to calculate the Mono/Poly ratios are shaded (see methods). *** *p* ≤ 0.001, Welch’s t-test. **(F)** Scheme showing cortical neurons grown on a microporous membrane enabling the separation of cell bodies and neurites for polysome profiling. Representative polysome profiles **(G and H)** and Mono/Poly ratios **(I)** of the cell body (blue) or neurite layer (purple) (n= 4). Areas measured to calculate the Mono/Poly ratios are shaded. * *p* ≤ 0.05, Welch’s t-test.

Polysome profiling is a biochemical fractionation method that allows one to examine the degree of ribosome association of a transcript, i.e. a monosome (single ribosome) or a polysome (integral ribosomes loaded on an mRNA) (11). Using polysome profiling, we examined the ribosome occupancy of transcripts in the hippocampus comparing area CA1 somata and neuropil that were microdissected from *ex vivo* rat hippocampal slices (Fig. 1B). We obtained a typical polysome profile with two ribosomal subunit peaks (40S and 60S), one monosome or single ribosome (80S) peak and multiple polysome peaks. We assessed the relative association of transcripts with monosomes or polysomes (M/P ratio) in the somata and neuropil by measuring the area under the curve of the corresponding absorbance peaks. While a large proportion of transcripts were associated with polysomes in the somata (Fig. 1C), the monosome to polysome ratio was greater than two-fold higher in the neuropil (Fig. 1D and E). To confirm the difference in the monosome to polysome ratios between somata and neuronal projections, we used a well-established *in vitro* system to enrich for cell bodies and neuronal processes (12). Neurons were grown on microporous membranes allowing dendrites and axons (but not cell bodies) to extend to the area beneath the membrane (Fig. 1F, fig. S1B and C). We separately harvested the cell bodies and dendrites/axons and again conducted polysome profiling. Consistent with the microdissected slice data, the monosome to polysome ratio was again significantly higher in neurites compared to cell bodies (Fig. 1, F to I).

### Monosomes actively elongate transcripts in the synaptic neuropil

In mammalian cells, polysomes are thought to represent the translationally active ribosome population (*13–15*). In contrast, monosomes, reflecting single ribosomes detected on transcripts, are presumed to represent the isolation of protein synthesis initiation and termination events, but not active protein synthesis (e.g. the elongation of the polypeptide chain). We compared the translational status of somatic or neuropil-localized monosomes and polysomes by precisely mapping the position of the ribosome(s) along the mRNA using ribosome profiling (16) (Fig. 2A). Monosomal or polysomal fractions from neuropil or somata were collected; the purity of fractionation was independently demonstrated by the lack of polysome or monosome peak on sucrose gradient profiles from isolated monosomal and polysomal fractions, respectively (fig. S2). Following polysome profiling, ribosomal fractions were digested and monosome or polysome footprint libraries were prepared. After sequencing three replicates of monosome/polysome footprint libraries and aligning the reads to a reference genome (alignment statistics shown in fig. S3A), the classical ribosome profiling quality metrics were assessed (fig. S3). As expected, the monosome and polysome footprints peaked at a length of around 31 nucleotides (representing the area occupied by the ribosome; fig. S3B and C) and exhibited a depletion of read densities in the untranslated regions (UTRs) and introns (fig. S3D and E). The ribosome profiling libraries were highly reproducible between replicates, with the majority of samples exhibiting Pearson correlation coefficients > 0.95 (fig. S3F).

**Fig. 2.**
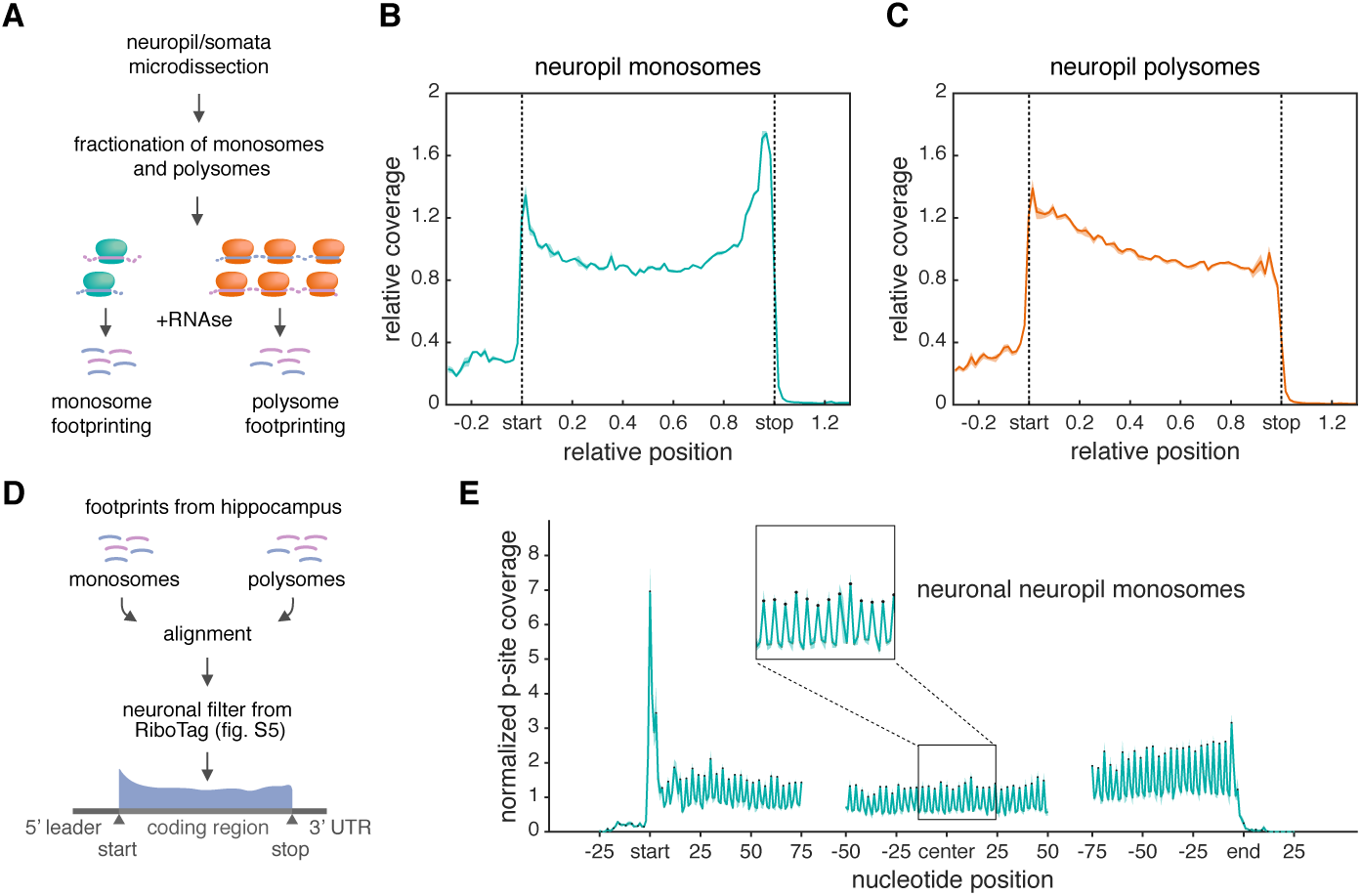
Neuronal monosomes actively elongate transcripts in the neuropil. **(A)** Experimental workflow. Somata or neuropil fractions were obtained, mono-/polysomes were isolated by polysome profiling and then ribosome profiling was performed on isolated fractions. **(B and C)** Metagene analyses showing the footprint density throughout the transcript open reading frame in the neuropil monosomes **(B)** or polysomes **(C)**. The average relative normalized coverage is plotted per nucleotide position, and the standard deviation is shaded (n= 3). Genes were individually normalized. **(D)** To assess the translational status of neuronal monosomes or polysomes, only reads classified as excitatory neuron-specific (see Fig. S5) were retained for further analysis. **(E)** Metagene analyses showing the P-site coverage of neuronal transcripts in the neuropil monosome sample. The average normalized coverage is plotted per nucleotide position around the 5’ end (start), central portion (center) and 3’end (stop) of the ORF. The standard deviation is shaded (n= 3).

We examined the positions of the RNA footprints obtained from neuropil monosomes (Fig. 2B) or polysomes (Fig. 2C) across the open reading frame (ORF) of transcripts. Both the monosome and polysome footprint coverage peaked at the 5’ ORF (near/at the translation initiation site); monosome footprints decreased more sharply than polysome footprints over the first 25% of the ORF before reaching a plateau. Only the monosome sample exhibited a pronounced enrichment of footprint reads around the stop codon, presumably reflecting the position of terminating ribosomes. This pattern is in good agreement with previously published metagene analyses of monosome and polysome footprint densities in yeast (17) thus confirming the purity of isolated monosomal and polysomal fractions. Surprisingly, however, a large fraction of monosome footprints occupied the center of the ORF, demonstrating that the localized monosomes are engaged in peptide elongation. A similar pattern was evident for the monosome (and polysome) footprint coverage in the somata (fig. S4A and B). Comparable results were obtained when performing the same analysis in monosome and polysome footprint libraries from the whole (non-microdissected) hippocampus (fig. S4C and D), indicating that the observed read distribution profiles were not a result of altered polysome integrity during the microdissection procedure.

As the somata and neuropil do not only comprise neurons but also glia and interneurons (fig. S5A), we developed a strategy to investigate the translational status of monosomes and polysomes in principal hippocampal excitatory neurons. In particular, we identified the translatome (ribosome-associated mRNAs) of select hippocampal excitatory neuron populations by combining RiboTag immunoprecipitation (RiboTag-IP) (18) with RNA-sequencing (fig. S5B and C). Using differential expression analysis (19), we identified transcripts enriched in the RiboTag-IP from hippocampi of Camk2Cre::RiboTag mice (fig. S5D and E) or microdissected somata (fig. S5D and F) and neuropil (fig. S5D and G) of Wfs1Cre::RiboTag mice. Combining the three datasets, we obtained a comprehensive list of 5069 mRNAs (“neuronal” transcripts) selectively translated in cell bodies and processes of excitatory hippocampal neurons (fig. S5H). The relative enrichment and de-enrichment of neuronal and glia/interneuron-related genes, respectively, was validated using a previously published dataset (fig. S5I) (20). These data were used to obtain a filtered list of neuronal footprint reads in monosome or polysome libraries from the somata and neuropil (Fig. 2D). As observed above, a significant fraction of neuronal transcripts displayed coverage in the elongating portion of the ORF in the monosome and the polysome samples of both neuropil (fig. S6A and B) and somata (fig. S6C and D). Footprints from actively elongating ribosomes also display a 3-nucleotide periodicity, reflecting the characteristic codon by codon translocation of the ribosome on its mRNA (16). Notably, the neuropil-derived monosome (Fig. 2E) and polysome (fig. S6E) footprints exhibit 3-nucleotide phasing in the 5’ORF, center and 3’ORF. These data indicate that both monosomes and polysomes contribute to the active elongation of transcripts localized to neuronal processes.

### Neuropil monosomes predominate on synaptic transcripts

To measure the degree to which a neuropil-localized transcript is translated by monosomes or polysomes, we focused on ribosomes that were undergoing elongation but not initiation or termination using footprints aligned to the center of the ORF (see Methods) in the monosome and polysome footprint libraries. Using DESeq2 (19), we identified localized neuronal transcripts preferentially translated by either monosomes or polysomes. In the neuropil, we found 463 transcripts significantly enriched in the monosome-versus 372 transcripts enriched in the polysome fraction (Fig. 3A). When we examined the footprint pattern across individual genes, we identified transcripts that displayed increased monosome (e.g. *Kif1a*; Fig. 3B) or polysome (e.g. *Camk2a;* Fig. 3C) footprint coverage throughout the entire ORF. There was also a large proportion of transcripts (e.g. *Slc17a7*; Fig. 3D) which exhibited equal coverage in monosome and polysome footprint libraries.

**Fig. 3.**
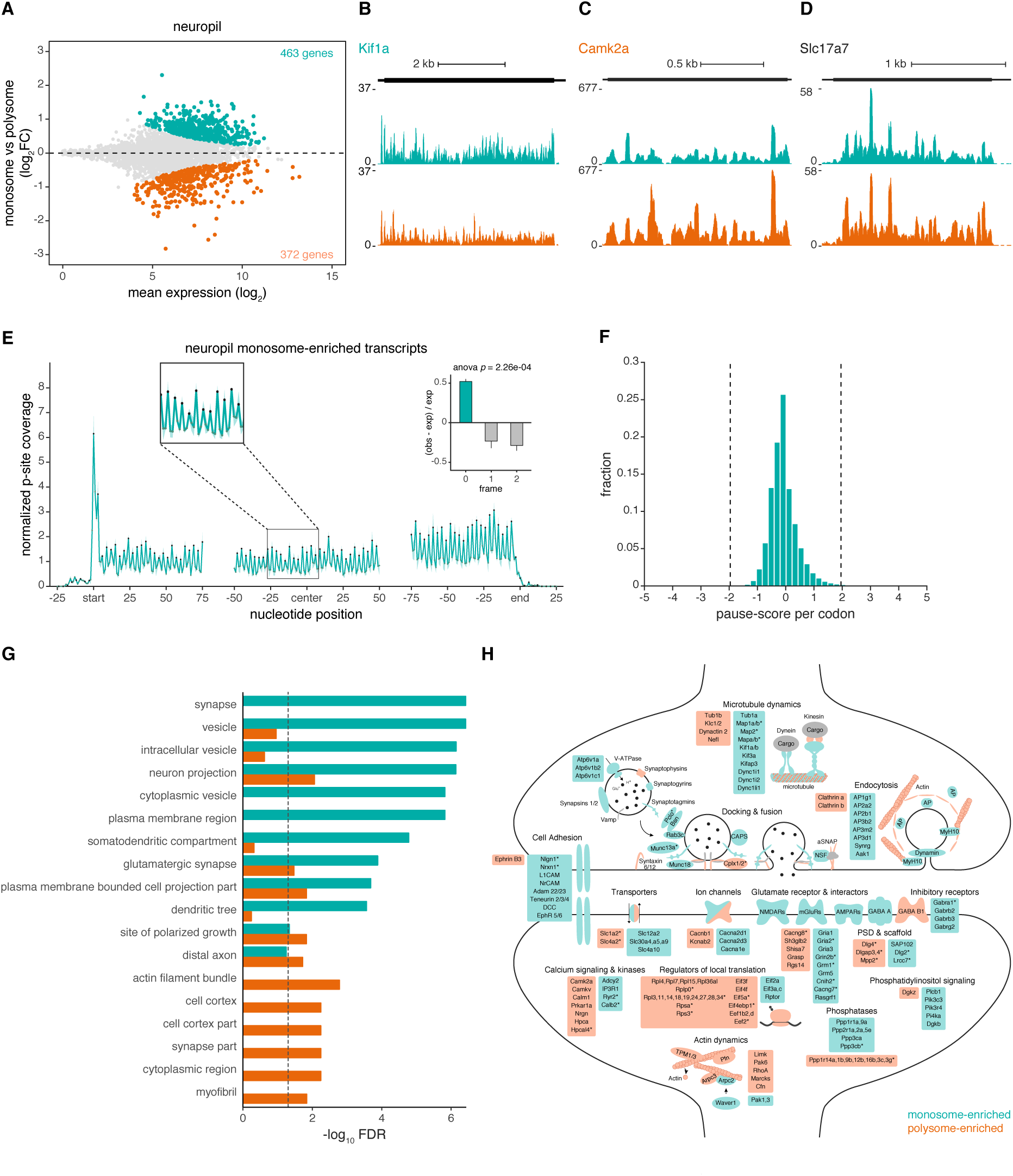
Local translation of key synaptic transcripts is predominantly accomplished by monosomes. **(A)** MA plot (the average, A, of the log read counts versus the differences in the log read counts, minus, M) showing transcripts with significantly enriched monosome (cyan) or polysome (orange) footprint coverage in the central portion of the ORF (region spanning 15 codons from the start site to 5 codons before the stop site). **(B to D)** Genome browser views representing the average monosome (top) or polysome (bottom) footprint coverage for 3 transcripts: *Kif1a* **(B)**, *Camk2a* **(C)** and *Slc17a7* **(D)**. Y axis indicates the number of normalized reads. **(E)** Metagene analysis showing the monosome P-site coverage of transcripts that exhibit significant monosome enrichment in the neuropil. The average normalized coverage is plotted per nucleotide position around the 5’ end (start), central portion (center) and 3’end (stop) of the ORF. The standard deviation is shaded (n= 3). Insets show the observed (obs) to expected (exp) ratio of the footprint distribution in different reading frames. *p* = 2.26e-04, ANOVA. **(F)** A pause score was computed for each codon located in the elongating ORF portion of the 463 monosome-enriched transcripts: pause score = normalized footprint coverage in monosome library – normalized footprint coverage in polysome library / (normalized footprint coverage in polysome library)^1/2^(n= 3). Fraction of codons per pause score. Dashed lines highlight pause score values of -/+ 1.96, values between these lines represent codons exhibiting similar coverage in monosome and polysome libraries. **(G)** GO terms representing the top ten significantly enriched protein function groups for monosome (cyan) or polysome (orange)-enriched transcripts. **(H)** Scheme of pre-and postsynaptic compartments highlighting some of the transcripts preferentially translated by monosomes (cyan) or polysomes (orange). Key synaptic components that were manually added owing to their exclusion by the excitatory neuron-specific filter are represented with an asterisk.

What transcript properties influence the neuropil monosome:polysome preference ? We detected a positive correlation between neuropil monosome to polysome preference and ORF length, 3’UTR folding energy or 5’UTR length (fig. S7A). On the other hand, an anticorrelation was found between the monosome:polysome ratio and the mean of the typical decoding rate index (MTDR, an estimate of the elongation efficiency (21)), GC-content, codon adaption index (CAI) or initiation rate (fig. S7A). We also observed an overrepresentation of uORF-containing transcripts (73 mRNAs) among monosome-enriched genes (463) (fig. S7B). Although a previous study in yeast reported that monosomes occupy non-sense mediated decay (NMD) targets (22), no relationship was found between the neuropil monosome to polysome preference of transcripts and their likelihood of classification as NMD targets (fig. S7C).

It has been proposed that a considerable fraction of synaptic mRNAs are associated with paused ribosomes (23, 24). Footprints from monosome-enriched transcripts exhibited a strong 3-nucleotide periodicity, suggesting that these mRNAs are not paused, but rather undergoing active elongation (Fig. 3E). Additionally, we assessed whether the increased footprint coverage of the monosome-preferring transcripts could result from pausing events at individual codons. Therefore, for the 463 monosome-enriched genes, we computed a pause score by comparing the normalized footprint coverage at individual codons between the monosome and polysome samples (see Methods). Most codons exhibited pause scores between −1.96 and +1.96 (Fig. 3F) indicating that, at these sites, there were no significant differences in pausing between the monosome and polysome libraries. Taken together, these findings demonstrate that monosome-enriched transcripts are actively elongated by single ribosomes in the neuropil.

To examine whether particular protein function groups are encoded by monosome-vs. polysome-preferring transcripts, we used gene ontology (GO) (Fig. 3G). We found that polysome-preferring transcripts often encode proteins involved in actin cytoskeleton remodeling (Fig. 3G and H). Because functional and morphological changes in synapses rely on the dynamic actin cytoskeleton remodeling (25), polysome-translation may be required to supply synapses with high copy numbers of cytoskeletal proteins. Interestingly, the monosome-preferring transcripts were enriched for GO terms such as ‘synapse’, ‘vesicle’ or ‘dendritic tree’, indicating that, in dendrites and axons, a significant proportion of transcripts important for synaptic function are principally translated by monosomes.

### Nature versus nurture of neuronal monosome-translation

We next assessed whether the localization of a transcript could affect its translational status. We noted with interest that, unlike the substantial monosome enrichment observed for neuropil transcripts (Fig 3A), a greater number of somatic transcripts exhibited a significant enrichment on polysomes (fig. S8A). To address whether the monosome to polysome preference is intrinsic to the transcript (nature) or influenced by the environment (nurture, i.e. the subcellular compartment), we compared the relative monosome to polysome enrichment between the neuropil and somata. We observed a high correlation (R^2^= 0.6) between somata and neuropil monosome to polysome ratios, indicating that a large proportion of transcripts prefer the same type of ribosome occupancy in both compartments (Fig. 4A monosome-enriched in quadrant 1 or polysome-enriched in quadrant 3). Indeed, 329 transcripts exhibited monosome preference in both somata and neuropil (fig. S8B). We also found, however, examples of transcripts that exhibited differential monosome-or polysome occupancy between somata and neuropil (e.g. monosome-enriched in one compartment and polysome-enriched in the other, or vice-versa, Fig. 4A quadrants 2 and 4). Notably, only a handful of transcripts exhibited a significantly lower monosome to polysome ratio in the neuropil than somata (Fig. 4A, purple dots). For example, *Arc*, an immediate early gene that regulates synaptic strength (26), is predominantly monosome-translated in the somata while its local translation preferentially involves polysomes (Fig. 4B). The majority of transcripts (e.g. *Serpini1* Fig. 4C) with differential ribosome occupancy between compartments displayed significantly elevated monosome to polysome fold-changes in the neuropil (Fig. 4A, cyan dots). Indeed, a cumulative frequency distribution indicated a significant shift towards higher monosome preference in the neuropil (Fig. 4D). Together, these results demonstrate that neuropil-localized transcripts are significantly more likely to be translated on monosomes than somatic transcripts.

**Fig. 4.**
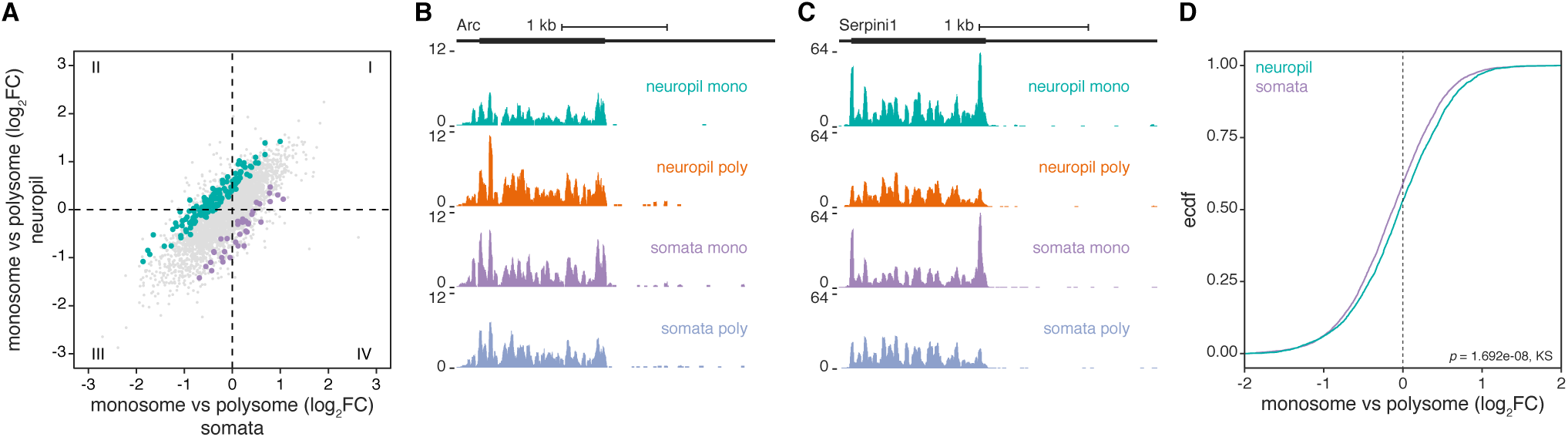
Localization influences the translational status of selective transcripts. **(A)** Monosome to polysome log2 fold-changes (FC) in the neuropil (y-axis) versus the somata (x-axis). The majority of transcripts exhibited correlated (R^2^= 0.6, *p* < 2.2e-16) monosome to polysome enrichments between both compartments. Colored dots highlight transcripts that exhibit significantly increased (cyan, n=136) or decreased (purple, n=36) monosome to polysome log2 fold-changes in the neuropil compared to the somata. Numbers represent the different quadrants. **(B and C)** Examples of transcripts that exhibited significant monosome-enrichment in the somata (*Arc*, **B**) or in the neuropil (*Serpini1*, **C**). **(D)** Cumulative distribution frequency depicting the monosome to polysome log2 fold-changes in the somata (purple) and neuropil (cyan) indicating a significant preference for monosome-mediated translation in the neuropil, *p* = 1.692e-08, Kolmogorov Smirnov test.

### Monosome translation contributes to the neuropil proteome

Individual synapses are small independent information processing units, each endowed with its own complement of proteins, ranging in copy numbers from 10s to a thousand or so (*27-30*). To study the contribution of monosome-and polysome-translation to the local proteome composition, we conducted mass spectrometry of neuropil proteins (see Methods) and estimated the neuropil absolute protein abundances using the iBAQ algorithm (intensity-based absolute quantification) (31, 32) (fig. S9). As might be expected, we observed higher median iBAQ values for proteins encoded by polysome-preferring transcripts when compared to proteins encoded by monosome-preferring transcripts (Fig. 5A). Paradoxically, though, when we plotted the neuropil monosome to polysome ratios of all transcripts over their respective protein abundance, we observed a surprisingly weak correlation (R^2^ = 0.021; *p*-value = 2.944e-11). Around half of the 326 proteins encoded by monosome-preferring transcripts exhibited protein abundances greater than average. (Fig. 5B), indicating that monosome-preferring transcripts can encode highly abundant proteins. In agreement with this, we found a weak relationship between neuropil monosome to polysome preference and previously published protein copy numbers in the pre-(Fig. 5C, (30)) and post-synapse (Fig. 5D, (27)). To further examine the properties of the low-versus high-abundance proteins encoded by monosome-preferring transcripts we divided the transcripts into “mono-low” (n=149) and “mono-high” (n=177) groups. Perhaps predictably, the “mono-high” transcripts exhibited greater mRNA levels (Fig. 5E and fig. S9B) and higher protein synthesis rates per mRNA (translational efficiency (TE); footprint over mRNA ratio) (Fig. 5F) in the neuropil when compared to the “mono-low” group. Taken together, our data highlight that predominantly monosome-translated transcripts contribute to the neuropil proteome composition through the encoding of a full range of low and high abundance proteins.

**Fig. 5.**
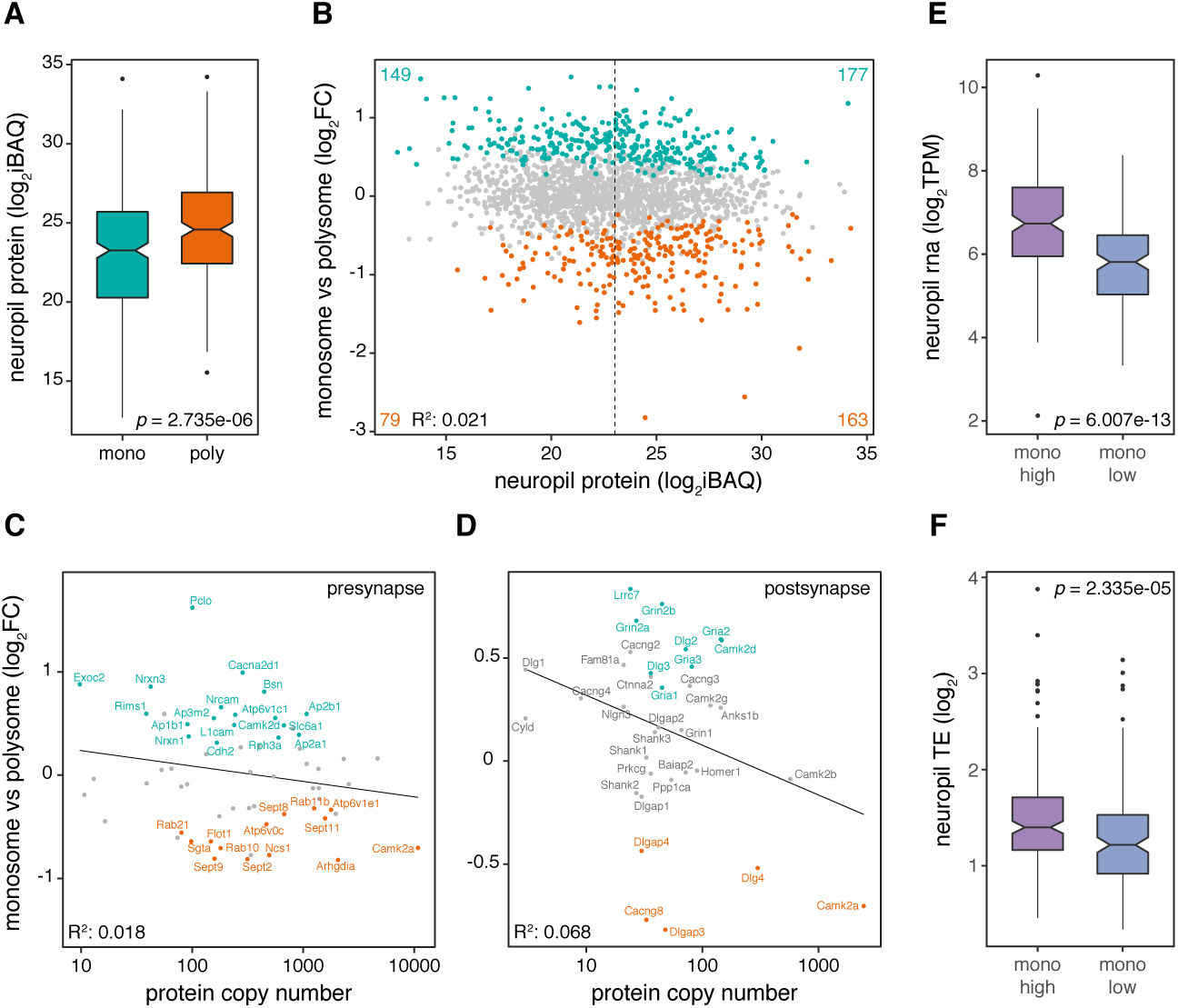
Monosome-preferring transcripts often encode abundant synaptic proteins. **(A)** Box plots of protein (log2 iBAQ) measurements in the neuropil for monosome-(mono, cyan, n=326) or polysome-(poly, orange, n= 242) enriched genes (Wilcoxon rank-sum test). **(B)** A scatter plot of the monosome to polysome fold-changes versus protein abundance (log2 iBAQ) for monosome (cyan)-, polysome (orange) -and non-enriched (grey) genes (R^2^ = 0.021, *p* = 2.944e-11). The dashed line indicates the mean log2 iBAQ value. **(C and D)** Monosome to polysome fold-changes in the neuropil were not correlated with the copy numbers of key pre-synaptic (Wilhelm et al. 2014) **(C)** and post-synaptic proteins (Loewenthal et al. 2015) **(D)**. Regression lines and corresponding adjusted Pearson’s R^2^ are represented (pre-synapse *p* = 0.1488, post-synapse *p* = 0.07145). **(E and F)** Box plots of RNA (log2TPM) (**E**) and translational efficiency (TE; footprint to mRNA ratio) (**F**) measurements in the neuropil for monosome-preferring genes encoding high-(mono-high, purple, n=177) or low-(mono-low, blue, n= 149) abundance proteins (Wilcoxon rank-sum test, RNA *p* = 6.007e-13, TE *p* = 2.335e-05).

## Discussion

In the present study, we investigated the translational landscape in neuronal processes and identified local translation on 80S monosomes as an essential source of synaptic proteins. To date, knowledge about the conformation of the translational machinery near synapses arises largely from electron micrographs where ribosomes are unambiguously identified when organized as a polyribosome cluster formed by more than three ribosomes (33). Monosomes, in contrast, are generally not quantified in EM, as their small size (10-25nm) does not allow one to distinguish them from other dark-staining particles in the cytoplasm (33). The sparse distribution of polysomes in dendrites and spines apparent in electron micrographs has led to the notion that local protein synthesis likely represents a minor source of synaptic protein in basal conditions (34). Indeed, until the recent detection of mRNAs and the machinery needed for their translation (*5, 35, 36*), the inability to identify polysomes in EM images from mature axons led to assertions that mature axons obtain protein via intracellular transport from the soma. We detected substantial levels of ongoing protein synthesis in the synaptic neuropil *in vivo* and identified a remarkably high number of both, pre-and post-synaptic transcripts that were preferentially translated on monosomes. This finding thus bridges the gap between the relative paucity of visualized translational machinery in neuronal processes and actual measurements of local translation.

The dendritic spines and their associated presynaptic boutons that comprise the excitatory synapse are independent cellular compartments that are very small, often below 100 nm^3^ for spines (37). Given this, the relatively large dimensions of a polysome (~100-200 nm (7)), poses limits for high occupancy in spines and axon terminals. Translation via smaller machines, i.e. monosomes, allows for a larger distribution of protein synthesis sites within synaptic compartments. While polysomes have been reported to move at an average speed of 2 µm/s (38), potentially greater dynamics of translating monosomes may allow them to patrol and serve a larger number of synapses. Each dendritic spine has been estimated to contain, on average, one polyribosome (6). Given that one polysome translates a single mRNA resulting in multiple copies of a single protein, this imposes constraints on both the timing and variety of locally synthesized proteins. We showed that neuropil-localized transcripts exhibit a greater monosome preference than somatic transcripts, potentially allowing for the production of a more diverse set of proteins from a limited pool of available ribosomes at synapses.

We found that monosome-preferring transcripts encode proteins that span a broad range of abundances in the neuropil. Because many synaptic proteins are present at very low copy numbers within the pre-and post-synaptic compartments (e.g. AMPARs; estimated to ~ 15-20 per PSD) (29), their local translation by single ribosomes may suffice to maintain or even alter the synaptic activity. We also uncovered a subset of monosome-preferring transcripts that encode surprisingly high-abundance proteins including the scaffolding proteins Bsn and Dlg3. The higher protein output of this group of transcripts is achieved, at least in part, by increased mRNA levels and more efficient translation. On the other hand, predominant polysome-translation was observed for key signaling, scaffolding or cytoskeletal proteins (e.g. Camk2a, PSD95, actin), which are present at very high copy numbers within synapses (29). Many studies investigating translational control in synaptic plasticity or neurological disorders have focused their analysis on transcripts that co-sediment with polysomes (11, 39–41). Given that monosomes are key contributors to the neuronal translatome, such analyses may provide only a conservative estimate of the translational regulation, especially when performed within synaptic compartments where single ribosomes likely substantially outnumber polysomes.

We showed that most transcripts exhibited a similar monosome to polysome-preference in both somata and neuropil, suggesting that ribosome occupancy is often an intrinsic feature of the transcript. We detected a positive correlation between the monosome:polysome ratio and ORF length, which is in agreement with previous studies reporting decreased initiation efficiency and protein abundance for long ORFs (*42–46*). Notably, contrasting observations have been made in yeast, where monosomes preferentially occupy short ORFs (22), suggesting differences in the translation mode of lower versus higher eukaryotes. We also observed a negative correlation between the neuropil monosome:polysome preference and initiation rate as well as elongation efficiency (i.e. MTDR and CAI). Thus, consistent with our observation that very abundant synaptic proteins are mostly translated by multiple ribosomes, polysome-preferring transcripts are often subject to positive translational regulation.

Some transcripts exhibited a differential monosome to polysome preference between the somata and neuropil. Neurons differentially localize 5’ and/or 3’ UTR isoforms between sub-cellular compartments (2, 35, 47, 48). Because these *cis*-regulatory mRNA elements regulate initiation efficiency (49, 50), neurons may fine-tune their monosome to polysome preference through selective targeting of competitive UTR isoforms between compartments. Interestingly, we found that *Arc*, a previously reported natural NMD target that contains 3’UTR introns (51), is monosome-preferring in the somata but polysome-preferring in the neuropil. According to the model proposed by Giorgi et al. (51), *Arc* may be silenced by NMD in the somata whereas, in the neuropil, synaptic activity could trigger its release from NMD resulting in a translational upregulation (i.e. polysome-translation).

Alternatively, differences in the monosome preference between somata and neuropil could also arise from differential localization/activity of specific translational regulators, including RNA-binding proteins (RBPs) (52, 53), microRNAs (54, 55), initiation/elongation factors (24, 41, 56, 57) or the ribosome itself (58). For instance, the RBP FMRP is thought to inhibit the translation of selective transcripts in neuronal processes by binding polyribosomes and pausing their translocation (59, 60). Synaptic activity has been reported to regulate the local translational machinery through changes in the phosphorylation status of initiation (57) and elongation factors (61). Thus, local activity-induced signaling events can also likely control the flow of ribosomes on an mRNA and dictate its monosome to polysome-preference.

A rapid up-regulation in the number of polyribosomes has been observed in electron micrographs of dendritic shafts and spines after synaptic plasticity induction (7). Our data show that monosome translation is the preferred mode of protein synthesis in neuronal processes and presumably satisfies the local demands under basal conditions. The formation of polysomes, however, could be required to supply synapses with *de novo* plasticity-related proteins in response to stimulation. Additionally, given the spatial limitations within dendritic spines and axonal boutons, synaptic activity could also regulate monosome translation to diversify the local proteome with spatial and temporal precision.

## Acknowledgements

We thank D. Vogel for assistance with the preparation of cultured neurons, M. Heumüller, T. Dalmay and J.J. Letzkus for assistance with the intracerebroventricular injections; I. Wüllenweber and F. Rupprecht for assistance with the proteomics analysis; N.T. Ingolia and MJ McGlincy (Dept. of Molecular and Cellular Biology, University of California, Berkeley, California 94720) for advice on bioinformatic analysis of footprint libraries; E. Valjent (IGF, CNRS, INSERM, University of Montpellier, Montpellier France) for providing the Wfs1Cre transgenic mice.

## Author contributions

A.B. and C.G designed, conducted and analyzed experiments. G.T. analyzed experiments. E.C. conducted experiments. J.D. Langer acquired the proteomics data. E.M.S. designed experiments and supervised the project, A.B. and E.M.S. wrote the manuscript. All authors edited the paper.

## Data and material availability

All data are available in the main text or the supplementary materials. The accession number for the raw sequencing data reported in this paper is: NCBI BioProject: PRJNA550323.

All proteomics data associated with this manuscript have been uploaded to the PRIDE online repository.

## List of supplementary materials

Materials and Methods

Fig. S1-S9

