## Supplemental Material for "Monosomes actively translate synaptic mRNAs in neuronal processes"

**Affiliations:**
[1]Max Planck Institute for Brain Research, Frankfurt, Germany
[2]Max Planck Institute of Biophysics, Frankfurt, Germany

[†] contributed equally
* to whom correspondence should be addressed:


**This PDF includes:**
Materials and Methods
Figs. S1 to S9

### Materials and Methods

#### 1. Experimental Procedures

##### Animals

Homozygous RiboTag Rpl22$^{HA/HA}$ mice (The Jackson Laboratory, 011029) were crossed with Camk2Cre (The Jackson Laboratory, 005359) or Wfs1Cre mice (The Jackson Laboratory, 009103). Eight-week-old C57Bl/6, Wfs1Cre::RiboTag and Camk2Cre::RiboTag mice were housed in standard cages and fed standard lab chow and water *ad libitum*. Wfs1Cre::RiboTag mice were treated for 3 days with tamoxifen (100 mg/kg, i.p., Sigma), dissolved in sunflower oil/ethanol (10:1) to a final concentration of 10 mg/ml, and used 1 week later for immunostaining or immunoprecipitation studies (*62*).

Adult male four-week-old Sprague Dawley SPF (Specific-Pathogen Free) rats from Charles River Laboratories were housed on a 12/12 hour light dark cycle with food and water *ad libitum* until sacrifice. Pregnant SPF females from timed matings from Charles River Laboratories were housed in the institute's animal facility for one week on a 12/12 hour light dark cycle with food and water *ad libitum* until the litter was born. Cortical neurons were derived from P0 (postnatal day 0) Sprague-Dawley rat pups (CD® Crl:CD, both male and female, RRID: RGD_734476). Pups were killed by decapitation.

The housing and sacrificing procedures involving animal treatment and care were conducted in conformity with the institutional guidelines that are in compliance with national and international laws and policies (DIRECTIVE 2010/63/EU; German animal welfare law; FELASA guidelines). The animals were euthanized according to annex 2 of § 2 Abs. 2 Tierschutz-Versuchstier-Verordnung. Animal numbers were reported to the local authority (Regierungspräsidium Darmstadt, approval numbers: V54-19c20/15-F126/1020 and V54-19c20/15-F126/1023).

##### Hippocampal tissue collection and microdissection

After sacrifice, the heads of four-week-old male rats or eight-week-old mice were immediately immersed in liquid nitrogen for 4s. The brains were removed and the hippocampi were rapidly dissected on an ice-cooled disk. Hippocampal slices (500 µm) were prepared as previously described (*63*). Tissue pieces were snap-frozen and kept at -80°C until lysis.

##### Primary cortical cultures

Dissociated rat cortical neurons were prepared from P0 day-old rat pups as previously described (*64*). Neurons were plated at a density of 100.000 cells/cm$^2$ onto poly-d-lysine-coated 100 mm, 3 µm pore polycarbonate membrane culture inserts (Corning 3420). At one DIV, AraC was added at a final concentration of 3 µM. After 2 days, medium was exchanged to pre-conditioned growth medium and neurons were cultured until 21 DIV. All cultures were maintained in a humidified incubator at 37°C and 5% $CO_2$. The sex of animals from which the cells were obtained was not determined.

### Immunolabeling of cortical neurons cultured on membrane inserts

At 21 DIV a part of the membrane was excised, shortly submerged in PBS pH 7.5 and fixed for 20 min in PFA (4 % paraformaldehyde in PBS pH 7.5). Cells were permeabilized with 0.5% Triton X-100 in PBS pH 7.5 supplemented with 4% goat serum for 15 min and blocked with blocking buffer (4% goat serum in PBS pH 7.5) for 1 hr. Dendrites were stained using an anti-MAP2 antibody (SySy 188004, 1:1000) in blocking buffer overnight at 4°C. After washing the cells three times for 5 min in PBS pH 7.5, the secondary antibody (ThermoFisher A488 A-11073, 1:1000) was incubated in blocking buffer for 45 min at room temperature. Cells were washed three times for 5 min in PBS pH 7.5 with DAPI added to the second wash. Membranes were mounted on glass slides using Aqua Poly/Mount and imaged from the top (cell body fraction) or bottom (neurite fraction).

### Tagged ribosome immunoprecipitation

HA-tagged ribosome immunoprecipitation of hippocampi from Camk2Cre::RiboTag or somata/neuropil sections from Wfs1Cre::RiboTag mice was performed as described previously (*65, 66*) with slight modifications. Tissue sections were homogenized in a glass homogenizer containing ice-cold RiboTag lysis buffer (50 mM Tris pH 7.4, 100 mM KCl, 12 mM $MgCl_2$, 1% NP40, 1 mM DTT, 20 U/ml SUPERaseIN*RNAse inhibitor (Ambion), 200 U/ml RNAsin (Promega), 100 µg/ml cycloheximide, 10 U/ml TurboDNAse, protease inhibitor (Roche)). After triturating the lysate 10 times using a 23 Gauge syringe, samples were chilled on ice for 10 min and cleared by centrifugation at 16.100g for 10 min. Ten percent of the supernatant were kept as an input. HA-immunoprecipitation (IP) was performed by incubation of the remaining supernatant with 5 µl of anti-HA antibody (abcam Ab9110) over night at 4°C with gentle rotation. Incubation of the samples with magnetic beads (Dynabeads protein G, Invitrogen), washes and elution were performed according to (*65*). Total RNA was extracted from both the input and immunoprecipitated ribosome-mRNA complexes, using the RNeasy MinElute kit (Qiagen). RNA integrity was assessed using the Agilent RNA 6000 Pico kit.

### Lysate preparation for polysome and ribosome profiling

*Tissue*

Tissue samples were homogenized in polysome lysis buffer (20 mM Tris pH7.5, 150 mM NaCl, 5 mM $MgCl_2$, 24 U/ml TurboDNAse, 100 µg/ml cycloheximide, 1 mM DTT, 1% Triton X100 and protease inhibitor mixture (Roche)) (*67*) by douncing in a glass homogenizer. After triturating the lysate 10 times using a 23 Gauge syringe, samples were chilled on ice for 10 min and cleared by two centrifugations at 16,100g for 6 min.

*Neuronal Culture*

At 21 DIV cortical primary neurons were washed twice in ice-old PBS pH 7.5 supplemented with 100 µg/ml cycloheximide. Neurons were collected with a scraper in polysome lysis buffer supplemented with 8% glycerol. After triturating the lysates 10 times using a 23 Gauge syringe, samples were chilled on ice for 10 min and then cleared by centrifugation at 16,100g for 10 min.

### Polysome profiling

Samples were loaded onto 10-50% sucrose density gradients that were prepared w/v in the following gradient buffer: 20 mM Tris pH 7.5, 150 mM NaCl, 5 mM $MgCl_2$, 100 µg/ml cycloheximide, 1 mM DTT. For polysome profiling from neuronal cultures, the gradient buffer was supplemented with 8% glycerol. Gradients were centrifuged for 2h 45 at 36.000 r.p.m at 4°C in a SW41 Ti swing-out rotor. Polysome profiling was performed using a density gradient fractionation system (Brandel) with upward displacement and continuous monitoring at 254 nm using a UA-6 detector. The area under the curve (AUC) of individual absorbance peaks was quantified. A monosome to polysome ratio was calculated by relating the monosome AUC to the sum of the AUCs of all polysome peaks. Fractions of 125 µl corresponding to the monosome or the polysome peaks were collected and pooled.

### Monosome and polysome footprint isolation

For the entire hippocampus monosome and polysome footprinting, 3 replicates, each comprising the hippocampi from three rats, yielding ~150 µg RNA, were used. For the somata and neuropil monosome and polysome footprinting, three replicates each comprising a pool of microdissected tissue from 55 rats, yielding ~110 µg RNA, was used. For each replicate, microdissected tissue was lysed as described above and aliquots containing 20 or 10 µg of RNA were retained for total ribosome footprinting and total RNA sequencing, respectively. The remaining lysate was loaded onto 10-50% sucrose gradients and centrifuged as described above. To prevent masking of the ribosome peaks by myelin (*68, 69*), each replicate was loaded onto 2-3 gradients and monosome or polysome fractions from different gradients were pooled after polysome profiling. A volume of monosome / polysome fraction containing 10 µg (hippocampi) or 2-5 µg (somata / neuropil) of RNA was diluted with gradient buffer and digested with RNAse I (epicenter) rotating for 45 min at 24°C as described previously (*70*). Nuclease digestion reactions were promptly cooled, spun and 10 µl SUPERaseIN*RNAse inhibitor was added. Samples were then layered onto a 34% sucrose cushion, prepared w/v in gradient buffer supplemented with 20 U/µl of SUPERaseIN*RNAse inhibitor. 80S particles were pelleted by centrifugation in a SW55Ti rotor for 3h 30 at 55.000 r.p.m at 4°C.

### Total ribosome footprint isolation

Neuropil lysates from three biological replicates (see section 'monosome and

polysome footprint isolation') containing 20 µg of RNA were digested with RNAse I (epicenter) shaking for 45 min at 400 r.p.m at 24°C as described previously (*70*). Nuclease digestion reactions were promptly cooled, spun and 10 µl SUPERaseIN*RNAse inhibitor was added. Samples were then layered onto a 34% sucrose cushion, prepared w/v in gradient buffer supplemented with
20 U/µl of SUPERaseIN*RNAse inhibitor. 80S particles were pelleted by centrifugation in a SW55Ti rotor for 3h 30 at 55.000 r.p.m at 4°C.

### Ribosome footprint library preparation

Footprint libraries were prepared according to McGlincy and Ingolia (2017) (*70*). After amplification, the libraries were run on an 8% non-denaturating TBE gel, 160 bp products were isolated and characterized using the Agilent High Sensitivity DNA assay. Libraries were sequenced on an Illumina NextSeq500, using a single-end, 52 bp run.

### RNA isolation and library preparation

RNA was isolated from tissue lysates using the Direct-zol RNA micro Prep kit (Zymo). RNA integrity was assessed using the Agilent RNA 6000 Nano kit. Rat neuropil total RNA sequencing libraries were prepared starting from ~200 ng total RNA using the TruSeq stranded total RNA library prep gold kit (Illumina). For the input/RiboTag-IP samples from Camk2Cre::RiboTag hippocampi or Wfs1Cre::RiboTag somata and neuropil, mRNA sequencing libraries were prepared starting from ~100 ng total RNA using the TruSeq stranded mRNA library prep kit (Illumina). Libraries were sequenced on an Illumina NextSeq500, using a single-end, 75 bp run.

### Mass spectrometry data acquisition

Three replicates of rat neuropil were microdissected as described above. Tissue pieces were snap-frozen and kept at -80°C until lysis. Tissue pieces were lysed in 4% Chaps, 8M Urea, 0.2M Tris HCl, 1M NaCl. All samples were digested, reduced and alkylated based on a previously published FASP-protocol (*71*). Dried peptide pellets were stored at -20°C until LC-MS/MS analysis. Proteolytic digests were analysed via Nano-LC-MS/MS on an Ultimate 3000 nanoUPLC (Thermo Fisher Scientific, Bremen) coupled to a Orbitrap Fusion Lumos (Thermo Fisher Scientific, Bremen).
After dissolving the dried peptides in 20 µl 0.1% FA in 5% acetonitrile, samples were separated using an Acclaim pepmap C18 column (50cm x 75µm, particle size 2µm) after trapping on an Acclaim pepmap C18 pre-column (2cm x 75µm, particle size 3µm). Trapping was performed for 6mins with a flow rate of 6µl/min using a loading buffer (98/2 water/acetonitrile with 0.05% Triflouroacetic acid). Peptides were then eluted and separated on the analytical column at a flow rate of 300nl/min with the following gradient: from 4 to 33% B in 150 min, 33 to 48% B in 20min, 48 to 90% B in 1min, and constant 90% for 13mins (buffer A: 0.1% FA in water, buffer B 0.1% FA in

80/20 acetonitrile/water). All LC-MS-grade solvents were purchased from Honeywell/ Riedel del Häen.
Peptides eluting from the column were ionised online using a Nano Flex ESI-source and analysed with an Orbitrap Fusion Lumos mass spectrometer in data-dependent-mode. Survey scans were acquired over the mass range of 350 - 1400 m/z in the Orbitrap (maximum injection time 50s, AGC, automatic gain control, fixed at 2x10E5 and R = 120K) and sequence information was acquired by a "Top-Speed" method with a fixed cycle time of 2s for the survey and following MS/MS-scans. MS/MS-scans were performed on the most abundant precursors exhibiting a charge state from 2 to 5 with an intensity minimum of 5x10E3. Picked precursors were isolated in the quadrupole at 1.4 Da and fragmented using HCD at NCE (normalized collision energy) = 30%. For MS/MS an AGC of 10E4 and a maximum injection time of 300s was used. Resulting fragments were detected in the Ion Trap using the rapid scan mode. The dynamic exclusion was set to 30sec with a mass tolerance of 10ppm. All samples were measured in technical triplicates.

### Intracerebroventricular puromycin administration

Mice were anesthetized with isoflurane (induction: 4%, maintenance: 2%) in oxygen-enriched air (Oxymat 3, Weinmann, Hamburg, Germany) and fixed in a stereotaxic frame (Kopf Instruments, Tujunga, USA). Core body temperature was maintained at 37.5°C by a feed-back controlled heating pad (FHC, Bowdoinham, ME, USA). Analgesia was provided by local injection of ropivacain under the scalp (Naropin, AstraZeneca, Switzerland) and systemic injection of metamizol (100 mg/kg, i.p., Novalgin, Sanofi) and meloxicam (2 mg/kg, i.p., Metacam, Boehringer-Ingelheim, Ingelheim, Germany) (*72*). A stainless steel 22-gauge guide cannula (PlasticsOne, Roanoke, VA) was implanted vertically towards the left lateral ventricle (A/P = −0.22 mm; Lat. −1 mm; D/V −2 mm). Guide cannulas were fixed onto the skull with instant adhesive and dental cement bonded to stainless steel screws anchored to the skull. An obturator was inserted into each guide cannula and remained in place until the drug infusion when it was removed and replaced with an injector that extended 0.5 mm beyond the tip of the guide cannula. After surgery recovery, 3 µl of puromycin solution (9 mg/ml, 10% DMSO/90% saline) or vehicle were infused for 1 min into the cannula through polyethylene tubing using an infusion pump (Stoelting) (*73*). The protein synthesis inhibitor control received an infusion of 3 µl of anisomycin (25 µg/µl, initially dissolved in 3N HCl and brought to pH7.3 by addition of 3N NaOH) (*74, 75*). Thirty min after the anisomycin infusion, mice were infused with 3 µl of puromycin (9mg/ml) supplemented with 75 µg anisomycin. After drug infusions, the tubing remained in place for 1 extra minute to ensure proper delivery of the solution. All mice were previously handled to ensure proper immobility during intracerebroventricular administration. 10 min after puromycin infusion, mice were transcardially perfused as described below.

#### Immunolabeling of hippocampal slices

After anesthesia with isofluorane, mice were rapidly euthanized and transcardially perfused with 4% (w/v) paraformaldehyde in PBS pH 7.5. Brains were post-fixed over night in the same solution and stored at 4°C. 30 µm thick sections were cut with a vibratome (Leica) and stored at 4°C in PBS pH 7.5, until they were processed for immunofluorescence. Hippocampal sections were identified using a mouse brain atlas and sections comprising between −1.34 and −2.06 mm from bregma were included in the analysis. Hippocampal sections from Wfs1Cre::RiboTag and Camk2Cre::RiboTag mice were processed as follows: free-floating sections were rinsed three times for 10 min with PBS pH 7.5. After 15 min incubation in 0.2% (v/v) Triton X-100 in PBS pH 7.5, sections were rinsed in PBS pH 7.5 again and blocked for 1 h in a solution of 3% BSA in PBS pH 7.5. Finally, they were incubated for 72 h at 4°C in 1% BSA, 0.15% Triton X-100 with the anti-HA antibody (abcam Ab9110, 1:500). In vivo puromycylated brain slices were immunostained as described previously (*73, 76*). Briefly, sections were incubated for 20 min with coextraction/fixation buffer (50 mM Tris-HCl, pH 7.5, 5 mM $MgCl_2$, 25 mM KCl, protease inhibitor mixture (Roche), 0.015% digitonin (Wako Chemicals), and 3% PFA and post-fixed for 15 min with 3% PFA. After three rinses with PBS pH 7.5, sections were incubated for 72 h at 4°C with puromycin (Milipore MAB E343, 1:1000) and Wfs1 (Proteintech 11558-1-AP, 1:1000) antibodies in a solution containing 0.05% saponin, 10 mM glycine, and 5% fetal bovine serum in PBS pH 7.5. After primary antibody incubation, sections were rinsed three times for 10 min in PBS pH 7.5 and incubated overnight at 4°C with the secondary antibody (ThermoFisher A546 A-11030, A647 A-21245, 1:500). Sections were rinsed three times for 10 min in PBS pH 7.5 and mounted in Aqua-Poly/Mount.

### 2. Data Analyses

#### Proteomics data analysis

Raw data were processed using the Max Quant software version 1.6.2.2 (*77*). MS/MS- spectra were searched against the UniprotKB-database from Rattus norvegicus (36080 entries, downloaded on 21/12/2017) and additionally against a database containing common mass spec contaminations using the probabilistic based algorithm from the Andromeda search engine. The set of stringent constraints allowed only peptides with full tryptic specificity allowing N-terminal cleavage to proline and up to 2 missed cleavages. Carbamidomethylation of cysteine was set as fixed modification. Oxidation of methionine and acetylation of the protein N-terminus were set as variable modifications. Minimum peptide length was set to 7 amino acids. The first search was performed with 20ppm precursor tolerance for mass recalibration and the main search mass tolerance was set to 4.5ppm. The fragment mass tolerance was 0.5 Da and the “match between runs” option was enabled. Peptides and proteins were identified based on a 1% FDR with the use of a decoy strategy and only those protein groups which were identified with at least 1 unique peptide were used for further analysis.

All proteomics data associated with this manuscript have been uploaded to the PRIDE online repository (*78*).

### Proteomics post processing

The Perseus package v1.6.2.2 (*79*) was used for further bioinformatic analysis of the resulting expression data from MaxQuant. Before further processing, decoy and contaminant database hits as well as proteins only identified using modified peptides ("identified by site") were excluded. Additionally only those protein groups which were identified in at least 2 out of 3 technical replicates and in 2 out of 3 biological replicates were considered for further analysis.

### Footprint genome and transcriptome alignment

Adapters were removed with Cutadapt v1.15 (*80*) (--cut 1 –minimum-length 22 –discard-untrimmed –overlap 3 –e 0.2). An extended UMI was constructed from the two random nucleotides from the RT primer and the five random nucleotides from the linker and added to the description line using a custom perl script. Trimmed reads that aligned to rat ncRNA were removed using Bowtie2 v2.3.4.3 (*81*) (--very-sensitive). Remaining reads were aligned to the rat genome (rn6) with the split-aware aligner STAR v2.6.1a (*82*) (--twopassMode Basic –twopass1readsN -1 –seedSearchStartLmax 15 –outSJfilterOverhangMin 15 8 8 8 –outFilterMismatchNoverReadLmax 0.1). When required, STAR --quantMode was used to retrieve transcript coordinates. The STAR genome index was built using annotation downloaded from the UCSC table browser. PCR duplicates were suppressed using a custom perl script and alignments flagged as secondary alignment were filtered out. All analyses mentioned below, except the differential expression analysis, were performed using the transcriptome alignments.

### RNA genome alignment

Adapters and low quality nucleotides were removed with Cutadapt v1.15 (*80*) (--minimum-length 25 –nextseq-trim=20). Reads were aligned to the rat (rn6) or the mouse (mm10) genome with STAR v2.6.1a (*82*).

### Assigning footprint reads to genomic features

Genomic feature coordinates (CDS, 3'UTR, 5'UTR, intron) were downloaded from the UCSC table browser as BED files (*83*). Bedtools v2.26.0 (*84*) was used to first convert BAM files into the BED format and second to identify reads overlapping with the individual features.

### Counting and differential expression analysis

*Monosome to polysome ratios*

Counts per gene were calculated from reads mapped to the genome using featureCounts v1.6.3 (*85*). Only a single transcript isoform, with the highest possible APPRIS score (*86*), was considered per gene. Only footprint reads aligned to the central portion of the ORF, by convention 15 codons from the start until 5 codons before the stop codon, were counted (*70*). Raw counts were fed into DESeq2 (*87*) for differential expression analysis. LFC shrinkage was used to generate more accurate log2 fold-change estimates (*88*). In order to test if the monosome to polysome fold change differs across compartments, an interaction was added to the design formula. In this analysis unshrunken log2 fold-changes were used.

*RiboTag-IP to input ratios*
Counts per gene were calculated from reads mapped to the genome using featureCounts v1.6.3 (*85*). All transcript isoforms were considered. Raw counts were fed into DESeq2 and LFC shrinkage was used.

### Classification of neuronal genes

A classifier to identify excitatory neuron-enriched genes was developed. The union of genes with significantly enriched RiboTag-IP to input fold-changes (FDR threshold of 0.05 on the adjusted p-value and a 30% enrichment) was formed from the three RiboTag experiments (Hippocampus Camk2Cre::RiboTag, somata/neuropil Wfs1Cre::RiboTag).

### Classification of non-sense mediated decay (NMD) targets

Genes with the Ensembl biotype annotation 'nonsense_mediated_decay' and 'retained_intron' were classified as possible NMD targets.

### Translational efficiency calculations

Translational efficiency was computed from three biological replicates of neuropil total ribosome footprinting. The translational efficiency of a gene was calculated as the ratio of the normalized footprint density to the normalized RNA-seq read density (as previously described in (*67*). In short, the normalized footprint density of a gene was determined by dividing the number of footprint reads in the gene's CDS by its CDS length in kilobases. This value was then normalized to the total number of footprint reads mapping to any region of the gene. The normalized RNA-seq read density was computed just as the normalized footprint density. Only genes with a minimum of 10 raw reads in all footprints as well as all RNA-seq libraries were used for analysis.

### Integration of proteomic and transcriptomic data

Protein and RNA data were matched as described in (*89*). A protein centric view was taken. For each protein in the protein group the corresponding RNA measures in transcripts per million (TPM) were summed and the mean of the corresponding translational efficiencies (calculated as described above) or monosome to polysome log2 fold-change was determined. In a functional group at least half the genes had to be classified as 'neuronal' in order to pass the neuronal filter. A functional group was determined as ‘monosome-enriched or polysome-enriched’, if the majority of its transcripts were classified as ‘monosome-enriched or polysome-enriched’. In all other cases, the functional group was classified as ‘non-enriched’.

#### Metagene analysis

Metagene plots represent the accumulated footprint coverage over the length-normalized ORF. The normalized footprint coverage was generated for each gene (footprint coverage divided by the average codon coverage). Edge positions were defined relative to the ORF start and stop codons and divided into 100 bins. Each gene contributed with its average normalized footprint coverage per bin.

#### 3-nucleotide periodicity analysis

First, the P-site offset was defined for individual footprint lengths. For this, all reads spanning the ORF start were used and the most probable offset from the start and end of the read was defined for each length.
Second, the P-site position per read was determined based on its length and the previously defined offset. All P-site positions were projected for 100 nucleotides around the ORF start, stop and center. The P-site coverage of each gene was normalized to its average footprint coverage. The nucleotide coverage at frame positions 0, 1 and 2 were assessed. To determine if the observed frame fraction differed from the expected frame fraction, a one-way analysis of variance (ANOVA) was performed. A significant p-value rejects the null-hypothesis that all frames exhibit the expected P-site coverage.

#### Genome browser track visualization

Footprint coverage was visualized as custom tracks on the UCSC Genome Browser (*90*).
Footprint alignments were converted into BedGraph files (https://genome.ucsc.edu/goldenPath/help/bedgraph.html) using Bedtools v2.26.0.

#### Gene ontology analysis

GO enrichment of monosome- or polysome-preferring genes was performed using the R package clusterProfiler (*91*) with a Benjamini-Hochberg multiple testing adjustment and a false-discovery rate cut-off of 0.05, using all expressed genes in the neuropil as background. The simplify function with a cutoff of 0.7 was used to remove redundancy from enriched GO terms.

### Correlation between the monosome to polysome fold-change and transcript attributes

DNA sequences were extracted from the rat (rn6) version genome. Only genes with valid values for all transcript attributes were used for analysis. The length of 3'- and 5'UTRs was set to a minimum of 10 nts.

*GC content*
The GC content was assessed by counting the number of G or C bases in the sequence and then dividing by the number of bases in the predicted 5'UTR, CDS or 3′ UTR.

*Minimum free energy (MFE)*
The ViennaRNA package version 2.0 with RNAfold was used to calculate the minimum free energy per 5'UTR or 3′ UTR sequence (*92*). A method described by Trotta et al. (*93*) was adapted to normalize minimum free energy units to the sequence length. The sequence length was restricted to a maximum of 500 nts in proximity to the start and stop codon. Only MFE values between 0 and -1 were used for analysis.

*Codon adaptation index (CAI)*
CAI values in the neuropil were obtained for neuronal genes only following the procedure described in (*94*).

*Initiation rate*
The initiation rate per gene was calculated based on the neuropil total ribosome footprint and RNA coverages as previously described in (*95*). In short, the initiation rate depends on the translational efficiency (defined as described above), CDS length, average time for a ribosome to traverse the CDS and the normalized ribosome occupancy in the initial 10 codons of the CDS. The average elongation rate was assumed to be 4 codons / second (*96*).

*Mean typical decoding rate (MTDR)*
A per gene MTDR was calculated based on the neuropil total ribosome footprint coverage as previously described in (*97*). In short, each amino acid decoding time was defined as a convolution of an average decoding time (a gaussian component with the parameters $\mu$ and $\sigma$) and a pausing decoding time (an exponential component with the parameter $\lambda$). A model fitting procedure was used to deconvolve the two distributions and identify the three parameters per amino acid.
The geometric mean of all average decoding times ($\mu$) was calculated to determine the per-gene MTDR.

### Upstream open reading frame (uORF)

To identify transcripts containing uORFs, neuropil total ribosome footprint libraries from three replicates were used. Only genes with annotated 5'UTRs were considered. A string match algorithm was used to identify sequences within annotated 5'UTRs that are flanked by a canonical in-frame start and stop codon. Only sequences with a minimum length of 3 codons and at least 10 raw footprints in all three replicates were considered as uORFs.

### Codon pause score analysis

For each codon in neuropil monosome-enriched genes (CDS only), a pause score was calculated based on a z-score-like quantity: pause score = normalized footprint coverage in monosome library – normalized footprint coverage in polysome library / (normalized footprint coverage in polysome library)$^{1/2}$. The resulting distribution represents the number of codons per gene with different coverage (i.e. a measure of observed pausing).

### Ribotools

All tools used in this study are contained in one modular C++ program called ribotools. It relies on the HTSlib (https://github.com/samtools/htslib) for parsing BAM files. The source code and further notes on the algorithms can be found on our GitLab repository:
https://gitlab.mpcdf.mpg.de/mpibr/schu/ribotools

### Data and software availability

The accession number for the raw sequencing data reported in this paper is: NCBI BioProject: PRJNA550323
All proteomics data associated with this manuscript have been uploaded to the PRIDE online repository.

68. P. Bernabo *et al.*, In Vivo Translatome Profiling in Spinal Muscular Atrophy Reveals a Role for SMN Protein in Ribosome Biology. *Cell Rep* **21**, 953-965 (2017).
69. W. P. Lou, A. Baser, S. Klussmann, A. Martin-Villalba, In vivo interrogation of central nervous system translatome by polyribosome fractionation. *J Vis Exp*, (2014).
70. N. J. McGlincy, N. T. Ingolia, Transcriptome-wide measurement of translation by ribosome profiling. *Methods* **126**, 112-129 (2017).
71. J. R. Wisniewski, A. Zougman, N. Nagaraj, M. Mann, Universal sample preparation method for proteome analysis. *Nat Methods* **6**, 359-362 (2009).
72. E. Abs *et al.*, Learning-Related Plasticity in Dendrite-Targeting Layer 1 Interneurons. *Neuron* **100**, 684-699 e686 (2018).
73. A. Biever *et al.*, PKA-dependent phosphorylation of ribosomal protein S6 does not correlate with translation efficiency in striatonigral and striatopallidal medium-sized spiny neurons. *J Neurosci* **35**, 4113-4130 (2015).
74. A. E. Power, D. J. Berlau, J. L. McGaugh, O. Steward, Anisomycin infused into the hippocampus fails to block "reconsolidation" but impairs extinction: the role of re-exposure duration. *Learn Mem* **13**, 27-34 (2006).
75. J. Remaud *et al.*, Anisomycin injection in area CA3 of the hippocampus impairs both short-term and long-term memories of contextual fear. *Learn Mem* **21**, 311-315 (2014).
76. A. Bastide, J. W. Yewdell, A. David, The RiboPuromycylation Method (RPM): an Immunofluorescence Technique to Map Translation Sites at the Sub-cellular Level. *Bio Protoc* **8**, (2018).
77. J. Cox, M. Mann, MaxQuant enables high peptide identification rates, individualized p.p.b.-range mass accuracies and proteome-wide protein quantification. *Nat Biotechnol* **26**, 1367-1372 (2008).
78. J. A. Vizcaino *et al.*, 2016 update of the PRIDE database and its related tools. *Nucleic Acids Res* **44**, D447-456 (2016).
79. S. Tyanova *et al.*, The Perseus computational platform for comprehensive analysis of (prote)omics data. *Nat Methods* **13**, 731-740 (2016).
80. M. Martin, Cutadapt removes adapter sequences from high-throughput sequencing reads. *EMBnet.journal17.1*, (2011).
81. B. Langmead, S. L. Salzberg, Fast gapped-read alignment with Bowtie 2. *Nat Methods* **9**, 357-359 (2012).
82. A. Dobin *et al.*, STAR: ultrafast universal RNA-seq aligner. *Bioinformatics* **29**, 15-21 (2013).
83. D. Karolchik *et al.*, The UCSC Table Browser data retrieval tool. *Nucleic Acids Res* **32**, D493-496 (2004).
84. A. R. Quinlan, I. M. Hall, BEDTools: a flexible suite of utilities for comparing genomic features. *Bioinformatics* **26**, 841-842 (2010).
85. Y. Liao, G. K. Smyth, W. Shi, featureCounts: an efficient general purpose program for assigning sequence reads to genomic features. *Bioinformatics* **30**, 923-930 (2014).

86. J. M. Rodriguez *et al.*, APPRIS 2017: principal isoforms for multiple gene sets. *Nucleic Acids Res* **46**, D213-D217 (2018).
87. M. I. Love, W. Huber, S. Anders, Moderated estimation of fold change and dispersion for RNA-seq data with DESeq2. *Genome Biol* **15**, 550 (2014).
88. A. Zhu, J. G. Ibrahim, M. I. Love, Heavy-tailed prior distributions for sequence count data: removing the noise and preserving large differences. *Bioinformatics* **35**, 2084-2092 (2019).
89. J. Cox, M. Mann, 1D and 2D annotation enrichment: a statistical method integrating quantitative proteomics with complementary high-throughput data. *BMC Bioinformatics* **13 Suppl 16**, S12 (2012).
90. W. J. Kent *et al.*, The human genome browser at UCSC. *Genome Res* **12**, 996-1006 (2002).
91. G. Yu, L. G. Wang, Y. Han, Q. Y. He, clusterProfiler: an R package for comparing biological themes among gene clusters. *OMICS* **16**, 284-287 (2012).
92. R. Lorenz *et al.*, ViennaRNA Package 2.0. *Algorithms Mol Biol* **6**, 26 (2011).
93. E. Trotta, On the normalization of the minimum free energy of RNAs by sequence length. *PLoS One* **9**, e113380 (2014).
94. R. Jansen, H. J. Bussemaker, M. Gerstein, Revisiting the codon adaptation index from a whole-genome perspective: analyzing the relationship between gene expression and codon occurrence in yeast using a variety of models. *Nucleic Acids Res* **31**, 2242-2251 (2003).
95. A. K. Sharma *et al.*, A chemical kinetic basis for measuring translation initiation and elongation rates from ribosome profiling data. *PLoS Comput Biol* **15**, e1007070 (2019).
96. C. Wang, B. Han, R. Zhou, X. Zhuang, Real-Time Imaging of Translation on Single mRNA Transcripts in Live Cells. *Cell* **165**, 990-1001 (2016).
97. A. Dana, T. Tuller, The effect of tRNA levels on decoding times of mRNA codons. *Nucleic Acids Res* **42**, 9171-9181 (2014).

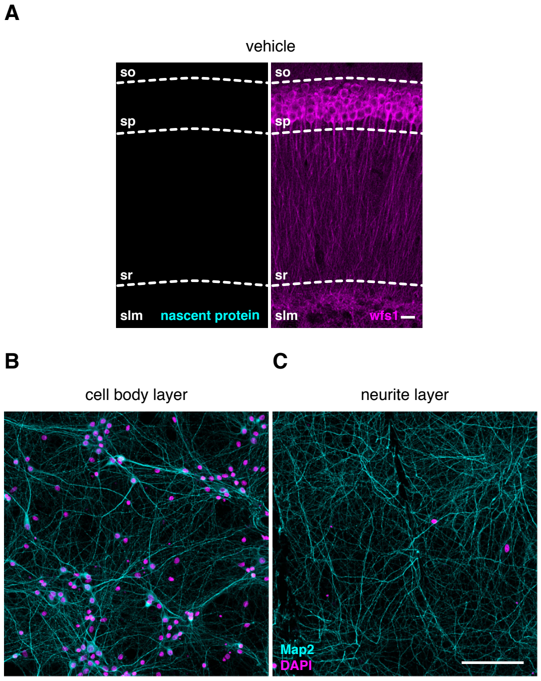


**Supplementary figure 1 related to figure 1.**

**(A)** Immunofluorescence staining of the nascent protein metabolic label puromycin (cyan) and the CA1 pyramidal neuron marker Wfs1 (purple) in hippocampal sections from control mice that received a brief infusion of vehicle into the lateral ventricle. Scale bar = 20 $\mu$m. so, *stratum oriens*; sp, *stratum pyramidale*; sr, *stratum radiatum*, slm, *stratum lacunosum moleculare*.

**(B and C)** Immunofluorescence staining of Map2 (cyan; dendrites) and DAPI (purple; nuclei) in the cell body **(B)** or neurite **(C)** fraction of cortical neurons grown on a microporous membrane. Scale bar = 100 $\mu$m.

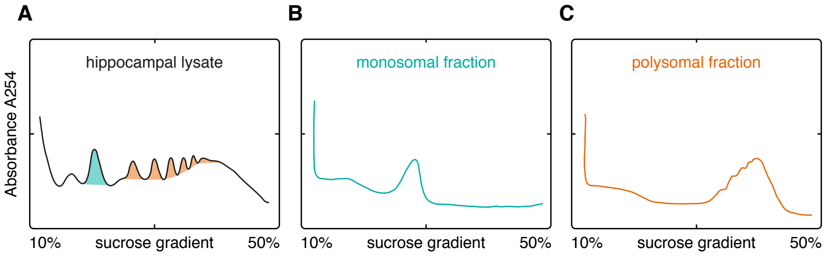


**Supplementary figure 2 related to figure 2.**
**(A to C)** Polysome profiling was performed on a hippocampal lysate **(A)**. The monosomal (cyan) and polysomal (orange) fractions were collected and then re-separated on a sucrose gradient. The purity of the isolated fractions is demonstrated by the lack of polysomes in the monosome sucrose gradient profile **(B)** or monosome in the polysome sucrose gradient profile **(C)**.

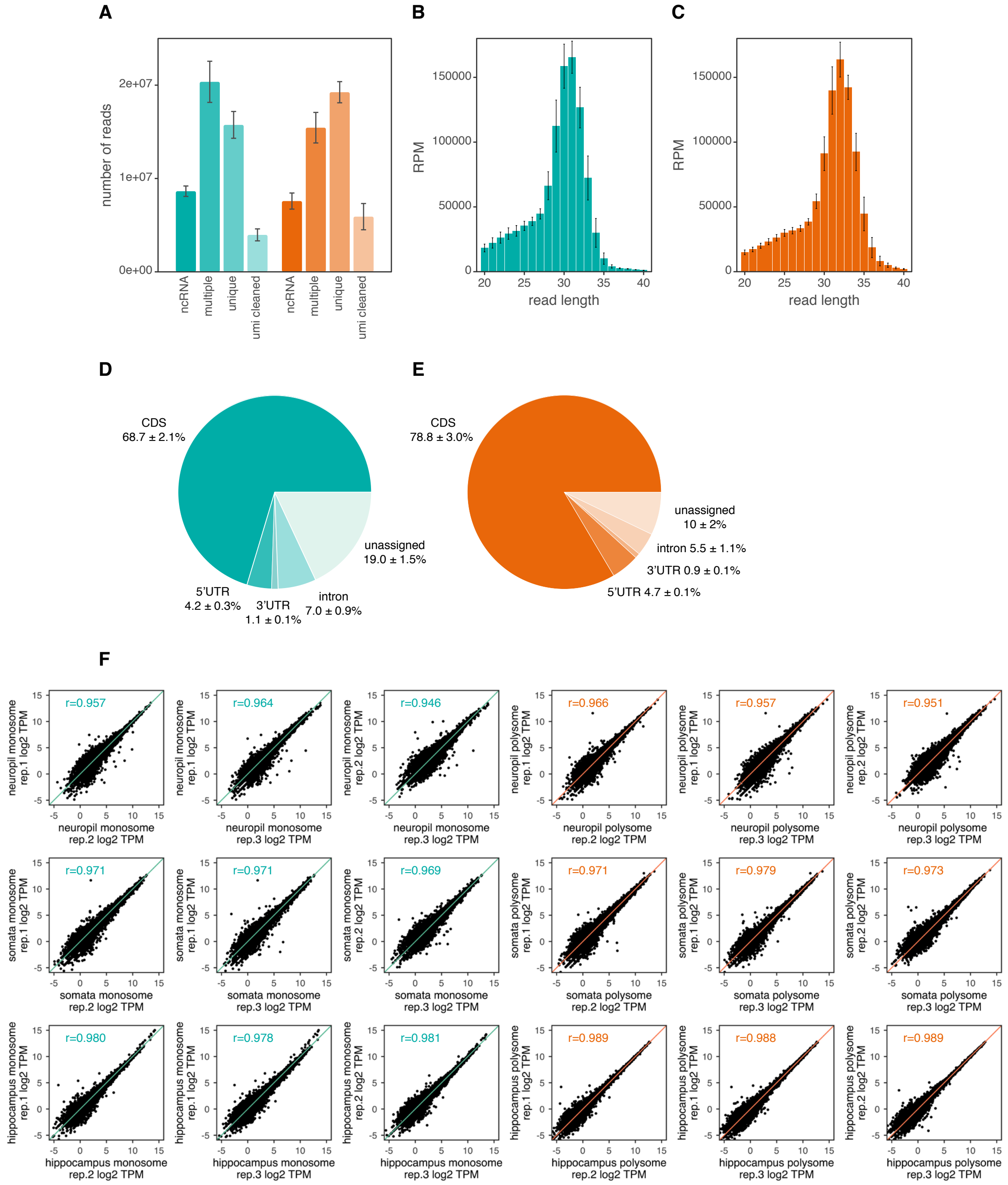


**Supplementary figure 3 related to figure 2.**

**(A)** Footprint read count by alignment fate (ncRNA, non-coding RNA; umi, unique molecular identifier; multiple, secondary alignment; unique, primary alignment) in all monosome (cyan) or polysome (orange) samples used in this study. **(B and C)** Distribution of footprint read lengths obtained in monosome **(B)** or polysome **(C)** libraries. **(D and E)** Fraction of monosome **(D)** or polysome **(E)** footprint reads aligning to various genomic features including the 5'UTR, CDS, 3'UTR and introns. **(F)** Scatter plots with Pearson's r correlation between biological replicates of hippocampus, somata and neuropil monosome/polysome footprint libraries.

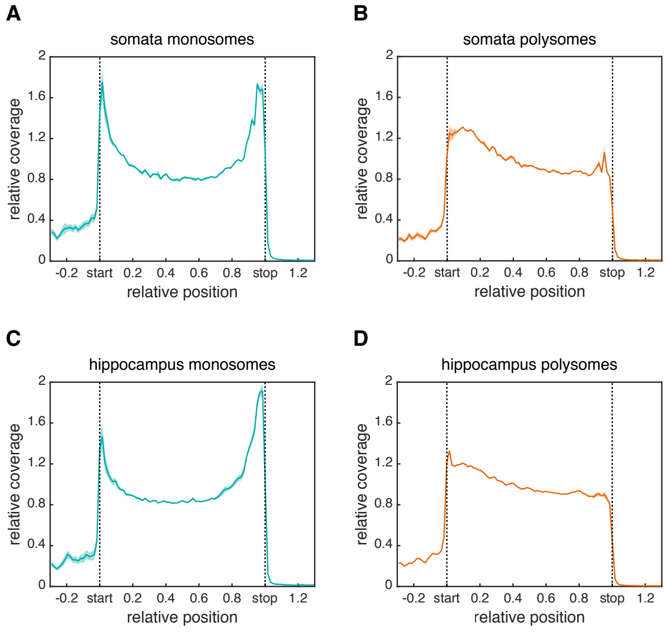


**Supplementary figure 4 related to figure 2.**

**(A to D)** Metagene analyses showing the footprint density in the monosome (cyan) or polysome (orange) samples from the somata **(A and B)** or hippocampus **(C and D)** throughout the open reading frame. The average relative normalized coverage is plotted per nucleotide position, and the standard deviation is shaded (n= 3). Genes were individually normalized.

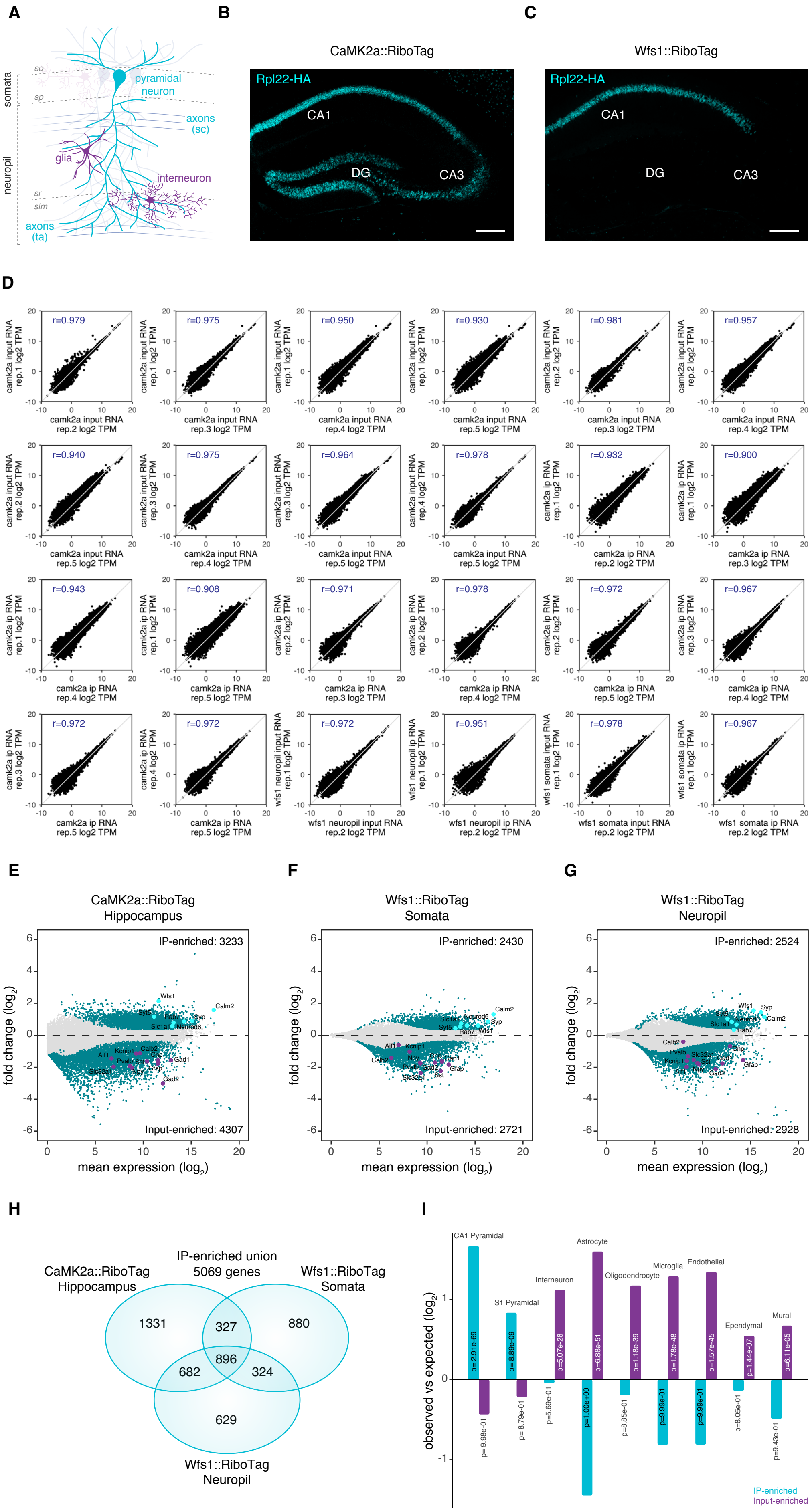
A
somata
neuropil
so
sp
sr
slm
pyramidal neuron
axons (sc)
glia
interneuron
axons (ta)
B
CaMK2a::RiboTag
Rpl22-HA
CA1
DG
CA3
C
Wfs1::RiboTag
Rpl22-HA
CA1
DG
CA3
D
r=0.979
camk2a input RNA rep.1 log2 TPM
camk2a input RNA rep.2 log2 TPM
r=0.975
camk2a input RNA rep.1 log2 TPM
camk2a input RNA rep.3 log2 TPM
r=0.950
camk2a input RNA rep.1 log2 TPM
camk2a input RNA rep.4 log2 TPM
r=0.930
camk2a input RNA rep.1 log2 TPM
camk2a input RNA rep.5 log2 TPM
r=0.981
camk2a input RNA rep.2 log2 TPM
camk2a input RNA rep.3 log2 TPM
r=0.957
camk2a input RNA rep.2 log2 TPM
camk2a input RNA rep.4 log2 TPM
r=0.940
camk2a input RNA rep.2 log2 TPM
camk2a input RNA rep.5 log2 TPM
r=0.975
camk2a input RNA rep.3 log2 TPM
camk2a input RNA rep.4 log2 TPM
r=0.964
camk2a input RNA rep.3 log2 TPM
camk2a input RNA rep.5 log2 TPM
r=0.978
camk2a input RNA rep.4 log2 TPM
camk2a input RNA rep.5 log2 TPM
r=0.932
camk2a ip RNA rep.1 log2 TPM
camk2a ip RNA rep.2 log2 TPM
r=0.900
camk2a ip RNA rep.1 log2 TPM
camk2a ip RNA rep.3 log2 TPM
r=0.943
camk2a ip RNA rep.1 log2 TPM
camk2a ip RNA rep.4 log2 TPM
r=0.908
camk2a ip RNA rep.1 log2 TPM
camk2a ip RNA rep.5 log2 TPM
r=0.971
camk2a ip RNA rep.2 log2 TPM
camk2a ip RNA rep.3 log2 TPM
r=0.978
camk2a ip RNA rep.2 log2 TPM
camk2a ip RNA rep.4 log2 TPM
r=0.972
camk2a ip RNA rep.2 log2 TPM
camk2a ip RNA rep.5 log2 TPM
r=0.967
camk2a ip RNA rep.3 log2 TPM
camk2a ip RNA rep.4 log2 TPM
r=0.972
camk2a ip RNA rep.3 log2 TPM
camk2a ip RNA rep.5 log2 TPM
r=0.972
camk2a ip RNA rep.4 log2 TPM
camk2a ip RNA rep.5 log2 TPM
r=0.972
wfs1 neuropil input RNA rep.1 log2 TPM
wfs1 neuropil input RNA rep.2 log2 TPM
r=0.951
wfs1 neuropil ip RNA rep.1 log2 TPM
wfs1 neuropil ip RNA rep.2 log2 TPM
r=0.978
wfs1 somata input RNA rep.1 log2 TPM
wfs1 somata input RNA rep.2 log2 TPM
r=0.967
wfs1 somata ip RNA rep.1 log2 TPM
wfs1 somata ip RNA rep.2 log2 TPM
E
CaMK2a::RiboTag
Hippocampus
IP-enriched: 3233
Input-enriched: 4307
fold change (log2)
mean expression (log2)
F
Wfs1::RiboTag
Somata
IP-enriched: 2430
Input-enriched: 2721
fold change (log2)
mean expression (log2)
G
Wfs1::RiboTag
Neuropil
IP-enriched: 2524
Input-enriched: 2928
fold change (log2)
mean expression (log2)
H
IP-enriched union
5069 genes
CaMK2a::RiboTag
Hippocampus
Wfs1::RiboTag
Somata
Wfs1::RiboTag
Neuropil
1331
327
880
896
682
324
629
I
observed vs expected (log2)
CA1 Pyramidal
S1 Pyramidal
Interneuron
Astrocyte
Oligodendrocyte
Microglia
Endothelial
Ependymal
Mural
IP-enriched
Input-enriched

**Supplementary figure 5 related to figure 2.**

**(A)** Scheme of the CA1 hippocampal area. The somata layer (sp) contains the cell bodies of CA1 pyramidal neurons (cyan). The neuropil layer (sr + slm) contains dendrites from pyramidal neurons (cyan) as well as axons (Schaffer collaterals (sc) + temporoammonic path (ta)). The somata and neuropil layers also contain interneurons and glia (purple). so, *stratum oriens*; sp, *stratum pyramidale*; sr, *stratum radiatum*, slm, *stratum lacunosum moleculare*. **(B and C)** Immunofluorescence staining of Rpl22-HA (anti-HA antibody; cyan) in hippocampal sections from Camk2aCre::RiboTag **(B)** and Wfs1Cre::RiboTag mice **(C)**. CA1, *cornu ammonis* 1; CA3, *cornu ammonis* 3; DG, *dentate gyrus*. Scale bar = 200 μm. (D) Scatter plots with Pearson's R correlation between biological replicates of whole hippocampi from Camk2aCre::RiboTag mice (n=5); and somata or neuropil from Wfs1Cre::RiboTag mice (n=2). **(E - G)** MA plots (the average, A, of the log read counts versus the differences in the log read counts, minus, M) showing differentially expressed transcripts between Rpl22-HA immunoprecipitation (IP) and input samples from: **(E)** whole hippocampi of Camk2aCre::RiboTag mice (n=5); **(F)** somata or **(G)** neuropil of Wfs1Cre::RiboTag mice (n=2). Cyan dots indicate the transcripts significantly enriched in the IP or input (DESeq2 with a threshold of 0.05 on the adjusted p-value and a 30%-fold-enrichment). Gray dots represent the non-enriched transcripts. Colored dots highlight markers for excitatory neurons (*Wfs1*, *Slc1a1*, *Calm2*, *Syp*, *Neurod6*, *Rab7* and *Syt5*); astrocytes (*Gfap*); microglia (*Aif1*); oligodendrocytes (*Cnp*) or different interneuron categories (*Gad1*, *Gad2*, *Kcnip1*, *Calb2*, *Npy*, *Pvalb*, *Slc32a1*, *Sst*). **(H)** Venn diagram comparing the transcripts significantly enriched in the Rpl22-HA IP from three different sources: the hippocampus of Camk2aCre::RiboTag mice, the microdissected somata or the neuropil of Wfs1Cre::RiboTag mice. A classifier to identify excitatory neuron-enriched genes was developed based on the union of the three datasets (5069 genes). **(I)** Observed-to-expected ratio of genes enriched in the Rpl22-HA IP (cyan) or the input (purple) and previously associated with different hippocampal cell-types by a single-cell study (*20*). Numbers within the bars indicate the p-values for the enrichment (using a hypergeometric test).

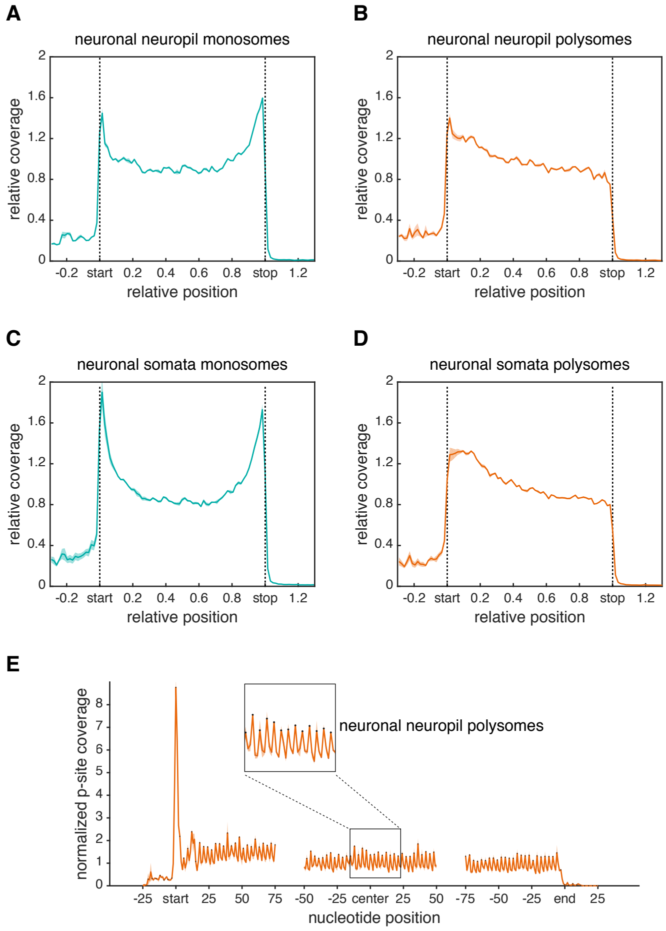


**Supplementary figure 6 related to figure 2.**
**(A-D)** Metagene analyses showing the footprint density across neuronal transcripts in the monosome (cyan) or polysome (orange) samples from the neuropil **(A and B)** and somata **(C and D)**. The average relative normalized coverage is plotted per nucleotide position, and the standard deviation is shaded (n= 3). Genes were individually normalized. **(E)** Metagene analyses showing the P-site coverage of neuronal transcripts in the neuropil polysome sample. The average normalized coverage is plotted per nucleotide position around the 5' end (start), central portion (center) and 3'end (stop) of the ORF. The standard deviation is shaded (n= 3).

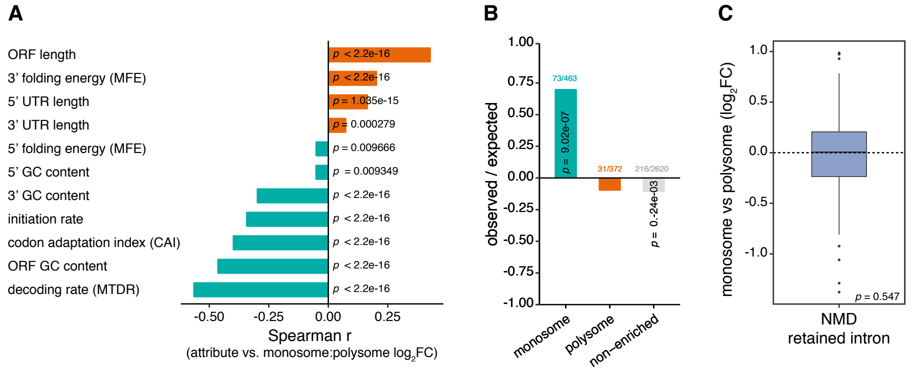


**Supplementary figure 7 related to figure 3.**
**(A)** Transcript attributes were correlated with the neuropil monosome to polysome fold-changes (FC). Positive (cyan) and negative (orange) Spearman correlation coefficients and p-values are shown. MFE, minimum free energy; CAI, codon adaptation index; MTDR, mean typical decoding rate. **(B)** Observed-to-expected ratio of monosome (cyan), polysome (orange)-and non-prefer-ring (grey) transcripts containing uORFs. Numbers of uORF-containing transcripts per gene subset are shown above the bars. Numbers within the bars indicate the significant p-values for over- and underrepresentation (using a hypergeometric test). **(C)** Box plot of monosome to polysome log2 fold-changes for transcripts classified as biotypes 'non-sense mediated decay (NMD)' or 'retained intron' by Ensembl (one-sample Wilcoxon signed rank test, $\mu = 0$).

**A**

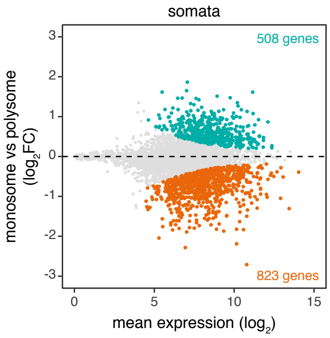


**B**

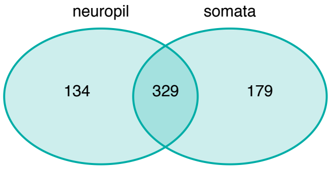


**Supplementary figure 8 related to figure 4.**

**(A)** MA plot (the average, A, of the log read counts versus the differences in the log read counts, minus, M) showing transcripts with differential monosome (cyan) or polysome (orange) footprint coverage in the central portion of the ORF (region spanning 15 codons from the start site to 5 codons before the stop site) in the somata. **(B)** Venn diagram representing the overlap of monosome-preferring transcripts between the somata and the neuropil.

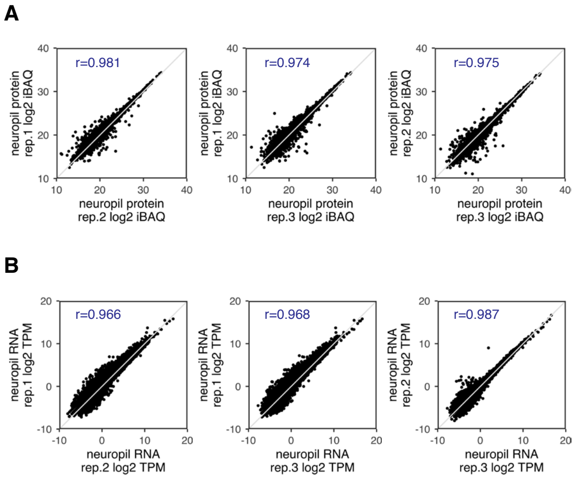


**Supplementary figure 9 related to figure 5.**
**(A and B)** Scatter plots with Pearson's R correlation of protein (log2-transformed iBAQ values) **(A)** and RNA (log2-transformed TPMs) **(B)** measurements from the neuropil of three biological replicates.
